# Interplay of hierarchical dynamics and their microscopic structures of polyampholyte gels and proteins

**DOI:** 10.1101/2024.04.26.591408

**Authors:** Yi Hui Zhao, Murugappan Muthukumar, Di Jia

## Abstract

Polyampholyte gel is a perfect physics model to mimic condensed state of proteins. We have studied the hierarchical dynamics of polyampholyte gels by dynamic light scattering. In addition to the normal gel mode, which indicates the gel elasticity, we also discovered a new mode with a stretched exponential decay with the stretched exponent β = 1/3, and a diffusive exponential decay, which indicates the coupled motion between counterion and the polyampholyte backbone. After dialysis to low salt concentration, the coupled motion of the counterion will go away, so that there are only two modes. Combined with a newly developed theory, we attribute this stretched exponential mode to hierarchical dynamics of the segments between two crosslinking junctions, whose segmental distribution obeys Poisson distribution. As salt concentration inside the gel increases, β decreases from 0.38 to 0.33, which is consistent with theoretical results. The gel with the molar charge ratio R=1, which is at the charge balance point, has the highest value β = 0.38. As long as R deviates further away from the charge balance point from either side, the β values decrease. When the gel is 100% positive charged, their dynamic light scattering results will go back to that of the normal polyelectrolyte gels.

## Introduction

Polyampholytes composed of oppositely charged functional groups are a physicochemical bridge between polymeric electrolytes and proteins^1,2^. They received much attention in the past 60 years due to their biocompatibility and similarity to biological systems such as proteins and intrinsically disordered proteins (IDPs)^3-6^. Polyampholyte chains and networks can undergo diverse conformational transformations including ordered, globular, coil, helix or stretched conformations^7-13^. Due to the conformational adaptability, the polyampholytes can reproduce some principal structural and thermodynamic features of biomacromolecules^14^. The conformation of these amphoteric substances resembles that of the proteins and IDPs, while their macromolecular structure relates them to the common polymer^5^. Thus synthesis polyampholytes are very important to duplicating the conformations, structures, functions and behaviors of proteins and IDPs^6,15^.

Lysozymes are naturally occurring enzymes present in a variety of biological organisms, such as bacteria, fungi, and animal bodily secretions and tissues. Lysozymes can act as an antimicrobial agent, an immune modulator, which stimulates the pro-inflammatory immune response and limits the inflammatory response^16-18^. The gelation of lysozymes and network junctions are formed by the association of two β-sheet fibrils through hydrophobic interactions^19,20^. The lysozyme gel exhibits antimicrobial, biodegradable, injectable and bio-adhesive properties^21^. The formation and dynamics of lysozyme amyloid fibrils and gels are key to investigate the functional structures of lysozyme and the association with protein deposition diseases^22,23^.

Polyampholyte hydrogels mimicking the solution properties of polyampholyte molecules have been prepared by copolymerization of ionic monomers possessing cationic and anionic charges in the presence of a chemical cross-linker^24-26^. The long-range Coulomb attraction between opposite charges causes certain limitations (frustrations) in the chain flexibility and the frustrations were shown to notably affect the dynamic behavior including swelling and phase transitions of the cross-linked polyampholyte hydrogels^27-30^. The frustration is a valuable instrument for understanding of folding mechanisms and organization of local functional structures in proteins^31,32^. The multiple relaxation dynamics of polyampholyte hydrogels is highly relevant in biomacromolecules, where charged macromolecules (proteins, IDPs) couple to their counterions and determine their three-dimensional structure, function, and ion mobility^33-35^. Studying the dynamic behavior of hydrogels with time can provide theoretical basis for the structural and functional changes caused by protein aging^36^. In this regard the dynamic behavior of synthetic polyampholyte hydrogels provide relevant models for elucidation of the structure function relations in biomacromolecules.

Polyampholyte hydrogels are soft materials with hierarchical structures, consisting of reversible ionic bonds at the 1-nm scale, a cross-linked polymer network at the 10-nm scale, and bicontinuous hard/soft phase networks at the 100-nm scale^37-39^. Sun synthesized self-healing polyampholyte hydrogels with high mechanical strength, good resilience because of the reversibility of the electrostatic and hydrogen-bonding interactions^40^. Gong et al. prepared high-toughness polyampholyte hydrogels by random copolymerization of oppositely charged ionic monomers which leads to the formation of multiple ionic bonds with a wide distribution of strengths. While the strong bonds act as permanent cross-links, the weak bonds acting as reversible sacrificial bonds easily fracture under a low stress and hence, dissipate energy to prevent crack propagation^38,41^. Many experimental studies of polyampholyte gels focused essentially on their swelling behavior^26,42,43^. Gong’s work shows that the shrinking and swelling kinetics of polyampholyte hydrogels could be intrinsically asymmetric and they calculated swelling and shrinking cooperative diffusion coefficients^44^.

The dynamic behavior of polymer gel systems are interesting candidates for dynamic-light-scattering (DLS) studies. In recent years, the polyelectrolyte gel modes and their contributions to both static and dynamic light scattering have been studied by many groups^45-51^. The time correlation function for polyelectrolyte gel can be fitted with a single-exponential function or a double-exponential function when the temperature is higher than the coil-to-globule transition temperature which due mainly to an appearance of microphase separation^50^. Svetlana Morozova prepared polyelectrolyte gel which correlation function can be fitted by a triple-exponential function, there is only one mode has a clear angle dependence which is the apparent elastic diffusion coefficient of the network^51^. For polyelectrolyte gels, the time correlation functions are all fitted by exponential decays, there is no stretched exponential decay. Recent dynamic light scattering (DLS) measurements on both dilute solutions of polyelectrolyte^52^ and trapped chains in gel matrix^53-55^ have shown a stretched exponential relaxation. The scattered intensity correlations of the lysozyme solution (10mg/ml) can be fitted with two modes^52^: one exponential decay and the other is stretched exponential decay which stretching parameter is 0.25 < β < 0.35. Jia reported a non-diffusive dynamical behavior of charged macromolecules embedded in a charged hydrogel^53^,displaying topologically frustrated localization driven by conformational entropy of the macromolecules. The dynamic light scattering (DLS) experiments show that the trapped molecules exhibit hierarchical segmental dynamics, and the time correlation function can be fitted by a combination of one or two exponential decays and one stretched exponential with stretched exponent β = 1/3. However, we report the first experimental evidence for β between a narrow range 1/3 < β < 3/8 for polyampholyte hydrogels, which is very different from polyelectrolyte gels and trapped chains in gel matrix. The hierarchical segmental dynamics and hierarchical structures of polyampholyte hydrogels are presented by addressing the coupling between the relaxations of polyampholyte networks, counterions from the polymer and added salt, and co-ions from the salt. The dynamic behavior of polyampholyte hydrogels can provide relevant models for elucidation of the structure function relations in proteins and IDPs.

## Experimental section

### Materials

Anionic monomer sodium 4-vinylbenzenesulfonate (NaSS), cationic monomer 3-(methacryloylamino) propyl-trimethylammonium chloride (MPTC) (50% weight/volume), UV initiator α-ketoglutaric acid were bought from Sigma-Aldrich, crosslinker bis-acrylamide solution (BIS) (2% weight/volume) and sodium chloride (NaCl) were purchased from Macklin. All materials were used without further purification. Deionized water was obtained from a Milli-Q water purification system (Millipore). The resistivity of deionized water used was 18.2MΩ cm. Hydrophilic polyvinylidene fluoride (PVDF) filters with 220nm pore were purchased from Millex Company.

### Synthesis of polyampholyte hydrogels

Polyampholyte physical hydrogels were synthesized using the one-step free-radical copolymerization. The gels were synthesized with different monomer concentration (Cm), salt concentration (Cs), crosslinking density and different molar charge ratio of negative charged monomers/ positive charged monomers (r). The crosslinking density of the gel is defined as the molar ratio of the cross-linker to the total amount of monomers. The procedure for the synthesis of the gel with Cm=2.0M, Cs=0M, r=1, 0.5 mol % crosslinking density and 0.1 mol % UV initiator (in relative to Cm) is an example. 3-(methacryloylamino) propyl-trimethylammonium chloride (MPTC) [(2.097mL) 50% (w/v)], 0.385mL 2% (w/v) of bis-acrylamide solution were put in a tube and Millipore water was added to bring a total volume of the solution to 5mL. Then sodium 4-vinylbenzenesulfonate (1.082g) and UV initiator α-ketoglutaric acid (1.5mg) was added to the tube. The pre-gel solution was bubbled with nitrogen gas for 30 min to remove any dissolved oxygen. The polymerization was irradiated with UV light for 12h. For the light scattering samples, the pre-gel solution was filtered (220nm PVDF filter) into a light scattering tube to remove the dust.

### Gel swelling

After polymerization, the as-prepared hydrogels were obtained. Swelling the as-prepared gel in pure water for one week to remove the counterions Na+ and Cl-, to tune the dynamics of ionic bonds, the as-prepared hydrogels were swelled in a series of NaCl solution with different concentrations for one week.

### Sample preparation and dynamic light scattering (DLS) measurement

Since DLS measurement is extremely sensitive to dust, the DLS tubes were first washed several times with pure water and acetone separately. After they were dried in the oven overnight, aluminum foil was used to wrap up the tubes and then these tubes were further cleaned by distilled acetone through an acetone fountain setup. Before polymerization, all the pre-gel solution were slowly filtered through 220nm PVDF hydrophilic filter into the tubes to remove the dust. All the sample preparation was conducted in a superclean bench to avoid dust.

DLS measurement was performed on a commercial spectrometer equipped with a multi-τ digital time correlator (ALV/LSE-5004) using a wavelength of 532nm laser light source. For each sample, the intensity at each of the scattering angles 30°, 40°, 50°, 60° and 70° was correlated, and the relaxation time averaged for three different spatial locations within the samples.

### Data acquisition and analysis

DLS measures the intensity-intensity time correlation function g_2_(q, t) by means of a multi-channel digital correlator and related to the normalized electric field correlation function g_1_(q, t) through the Siegert relation,

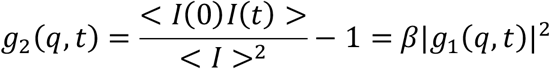

where β is the instrument coherence factor, t is the decay time and I(t) is the scattered intensity. The scattering wave vector q is defined by q=(4πn/λ)/sin(θ/2), where λ is the wavelength of the incident laser light (532nm), n is the refractive index of the solution (1.332), and θ is the scattering angle. The electric field-field correlation function g_1_(q, t) is fitted with a sum of several relaxation modes,

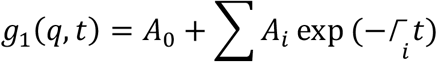

where A_0_ is the baseline, and A_i_ is the intensity weighting fraction of ith modes, each having a characteristic decay rate Г_i_. Both multiexponential fitting method and the CONTIN method were used to analyze the data. CONTIN analysis fits a weighted distribution of multiple relaxation decay times by the inverse Laplace transform. In the presence of multiple modes and in order to facilitate comparison to a theory, the normalized g_1_(q, t) was fitted using one or two exponential decays and a stretched exponential function. The decay rate Г_i_ is obtained from 2 for the multiexponential fits. If the mode is diffusive, then the decay rates are linear with q^2^ for all scattering angles. From the slope of Г = Dq^2^ across all q, diffusion coefficient D is obtained. The fitting residuals, which are obtained from the different between the raw data and the fitting curve, are randomly distributed near the mean of zero and do not have systematic fluctuations about their mean, indicating that the fitting quality is high. The fitting results obtained from both multiexponential fits and the CONTIN analysis are similar.

## Results and discussion

There are four parameters can be tuned, they are monomer concentration Cm, crosslinking density (XL), salt concentration Cs and molar charge ratio of negative charged monomers/ positive charged monomers (r).

### 1. Changing monomer concentration Cm before and after dialysis by deionized water

We have done DLS for two samples of monomer concentration Cm=2M, 2.5M, respectively with fixed crosslinking density 0.5% (labeled as 0.5XL), no additional salt (Cs=0 M), r=1 before dialysis (as prepared) in Fig. 1a-c and after dialysis by water in Fig. 1d-f. (Please note here Cs indicates additional NaCl salt concentration). Fig. 1g shows the β averaged values before and after dialysis.

**FIG. 1.**
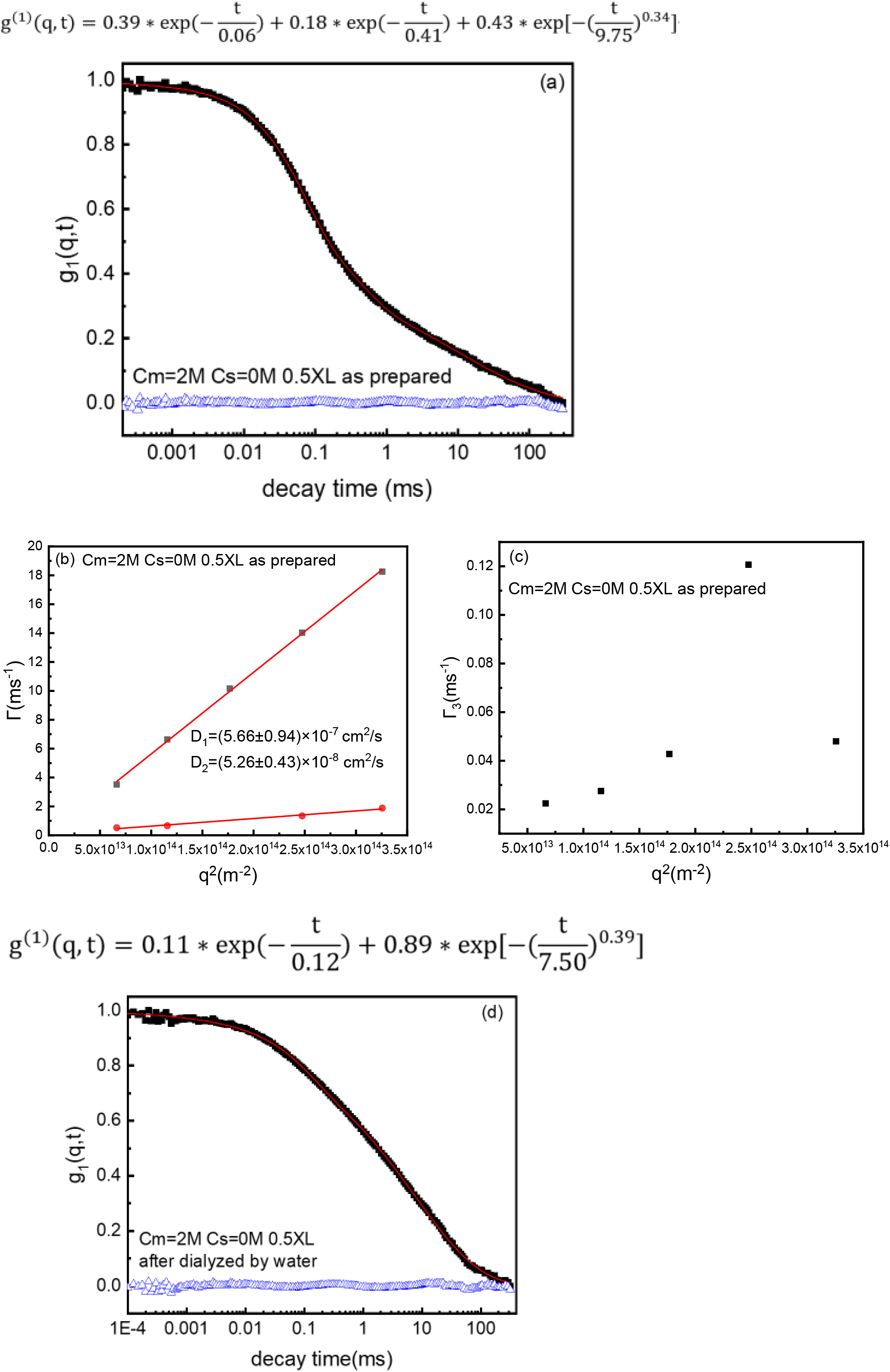
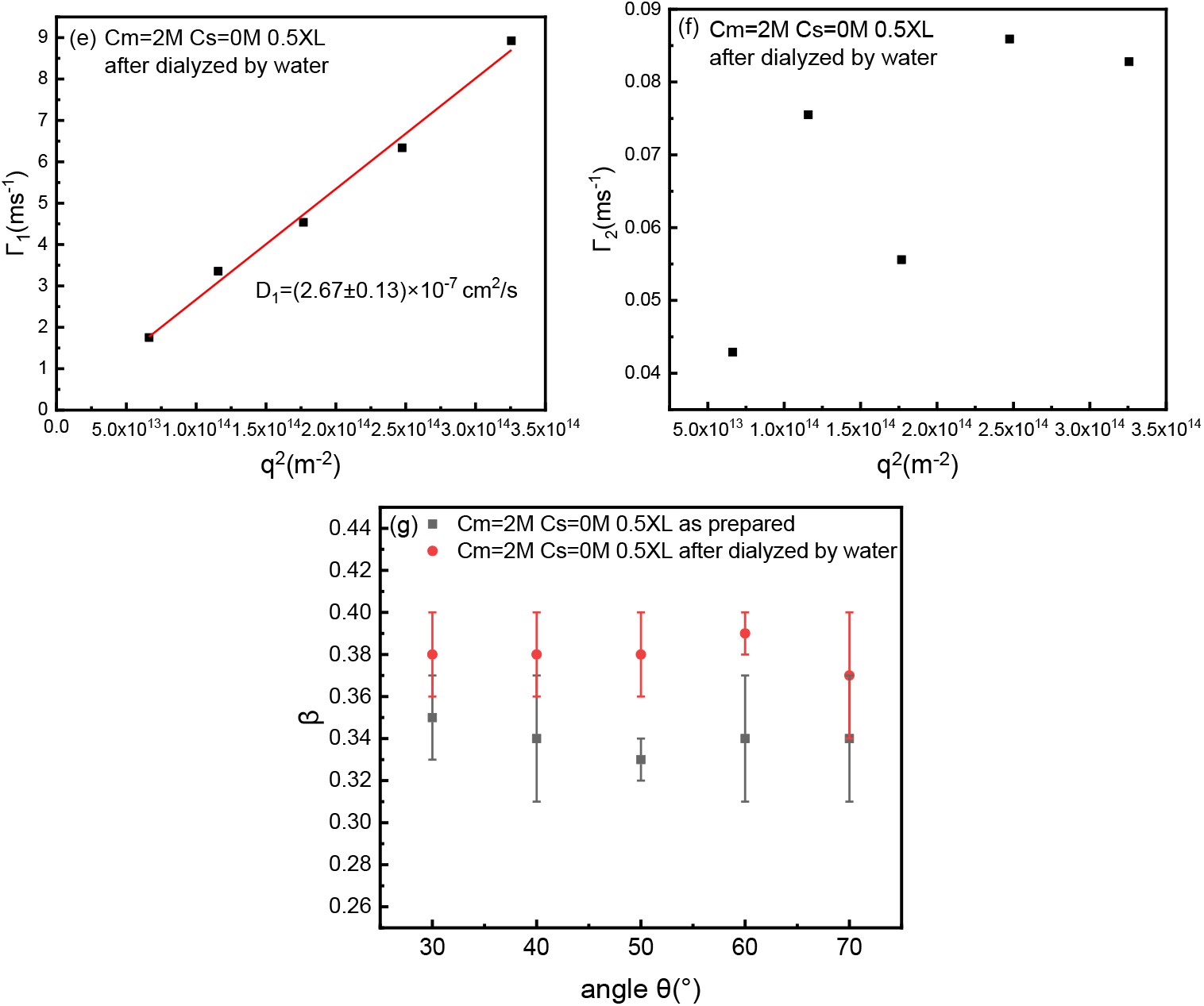
(a) Normalized field correlation function g_1_(t) at scattering angle 60° for as prepared gel at fixed Cm=2M, Cs=0M, r=1 and 0.5XL (red lines are the best fits and blue triangles are residuals). (b), (c) Corresponding q^2^ dependence of relaxation rates Г_1_, Г_2_, Г_3_ for as prepared gel at fixed Cm=2M, Cs=0M, r=1 and 0.5XL. (d) Normalized field correlation function g_1_(t) at scattering angle 60° for gels after dialyzed by water at fixed Cm=2M, Cs=0M, r=1 and 0.5XL (red lines are the best fits and blue triangles are residuals). (e), (f) Corresponding q^2^ dependence of relaxation rates Г_1_, Г_2_ for gels after dialyzed by water at fixed Cm=2M, Cs=0M, r=1 and 0.5XL. (g) Stretched exponent β of as prepared gel and gel after dialyzed by water. Table 1. Summary of changing Cm before and after dialysis for polyampholyte hydrogels.

### 2. Changing crosslinking density at fixed Cm=2M, and additional salt Cs=0 for the as-prepared gels

We have done DLS for samples with crosslinking density from 0.5% to 3.7%, with fixed monomer concentration Cm=2M, r=1, and no additional salt (Cs=0 M) before dialysis (as prepared samples).

All the gels with different crosslinking density can be fitted using three terms: two exponential decays and a stretched exponential decay with β around 1/3. The fitting function is as below:

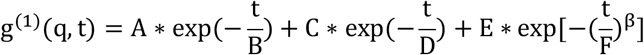

Both the first and second exponential decays are diffusive, so that we can get diffusion coefficients D_1_ and D_2_ respectively. As shown in Fig. 2, both D_1_ and D_2_ increases with crosslinking density. The stretched exponential term is non-diffusive and β is around 1/3.

**FIG. 2.**
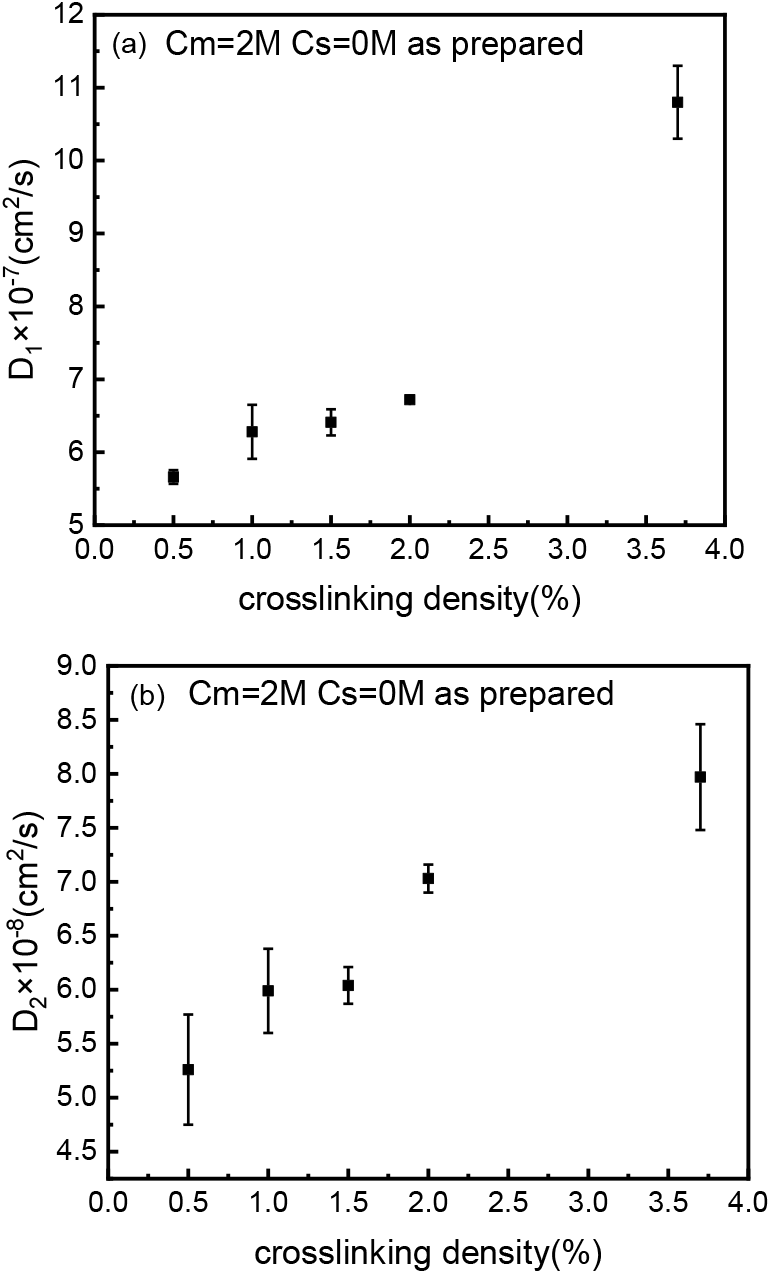
(a) Diffusion coefficients D_1_ for as prepared gels with different crosslinking density at fixed Cm=2M, r=1 and Cs=0M. (b) Diffusion coefficients D_2_ for as prepared gels with different crosslinking density at fixed Cm=2M, r=1 and Cs=0M.

### 3. Changing salt concentration (Cs)s at fixed Cm=2M and fixed crosslinking density 0.5% (labeled as 0.5XL), r=1 for gels before dialysis (as-prepared) and after dialysis

We synthesized the gels with Cm=2.0M and no additional NaCl salt. Then we do the dialysis for the as prepared gels by adding NaCl salt solutions with certain concentrations on top of the gel in the DLS tubes. We did the dialysis for 1 week, and we do the DLS measurements as a function of Cs of dialysis solution.

As shown in Table 3, as the Cs of dialysis solution. increases, the gel mode D_1_ increases, and β also decreases from 0.38 to 0.33, as shown in Fig. 3g-h and Table 3.

**FIG. 3.**
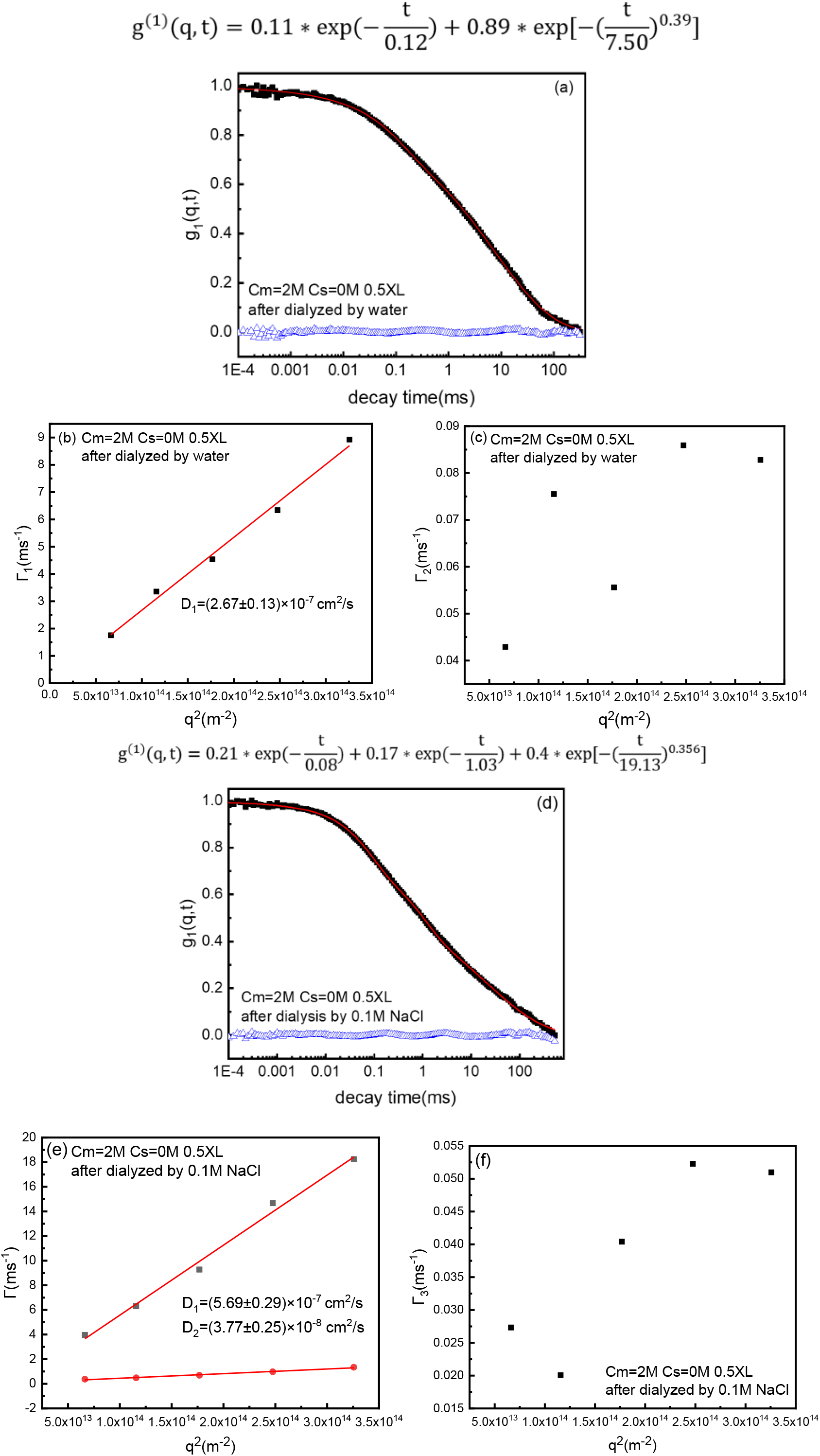
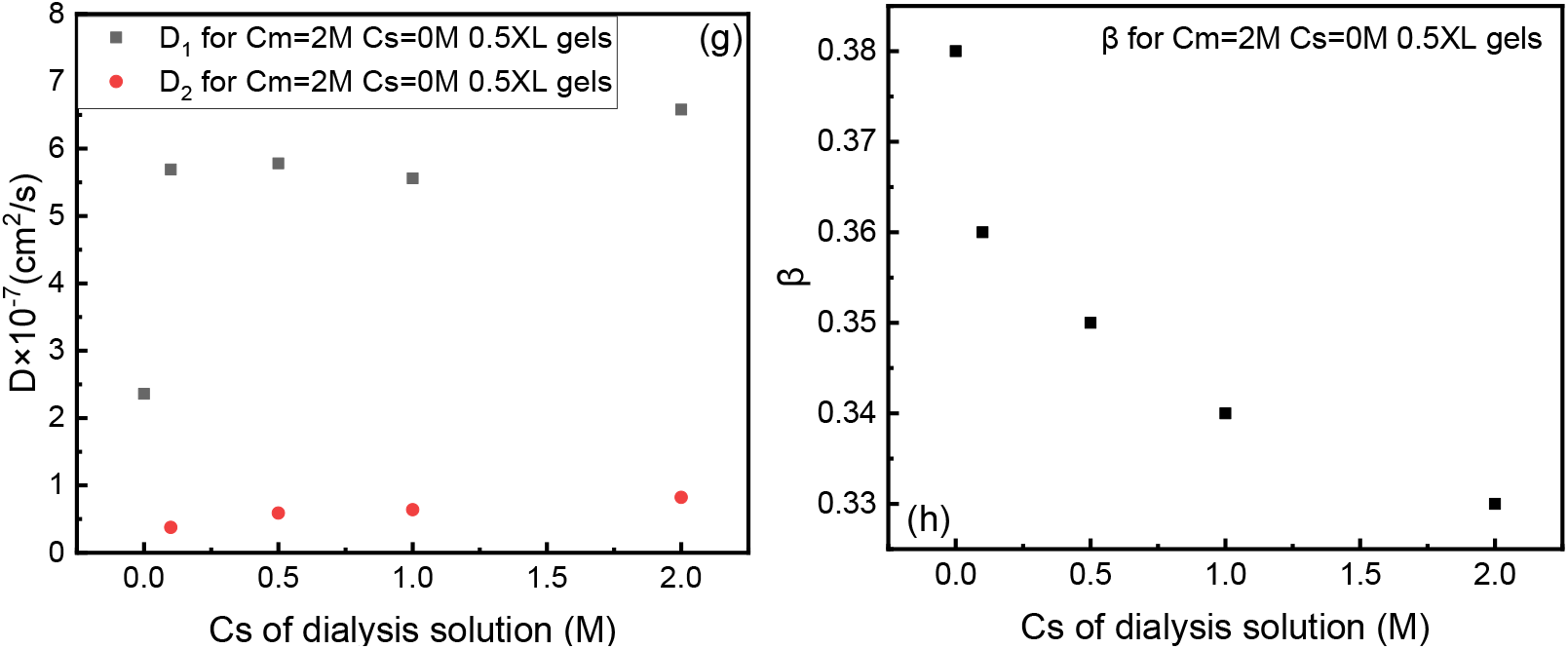
(a) Normalized field correlation function g_1_(t) at scattering angle 60° for as prepared gel at fixed Cm=2M, Cs=0M, r=1 and 0.5XL. (red lines are the best fits and blue triangles are residuals). (b), (c) Corresponding q^2^ dependence of relaxation rates Г_1_, Г_2_ for as prepared gel at fixed Cm=2M, Cs=0M, r=1 and 0.5XL. (d) Normalized field correlation function g_1_(t) at scattering angle 60° for gel at fixed Cm=2M, Cs=0M, r=1 and 0.5XL after dialyzed by 0.1M NaCl solution. (red lines are the best fits and blue triangles are residuals). (e), (f) Corresponding q^2^ dependence of relaxation rates Г_1_, Г_2,_ Г_3_ for gel at fixed Cm=2M, Cs=0M, r=1 and 0.5XL after dialyzed by 0.1M NaCl solution. (g) Diffusion coefficients D_1_ and D_2_ for Cm=2M, r=1, and 0.5XL gels at different Cs of dialysis solution. (h) Stretched exponent β of Cm=2M, Cs=0M, r=1, and 0.5XL gels at different Cs of dialysis solution.

### 4. Changing molar charge ratio r of negatively charged monomers/ positively charged monomers at fixed Cm=2M, Cs=0M and crosslinking density 0.5% (labeled as 0.5XL) for gels after dialysis by water

#### 4.1 gel with molar charge ratio r of negatively charged monomers/ positively charged monomers=3:7

at fixed Cm=2M, Cs=0M (no addition salt), 0.5% crosslinking density, dialysis by water. As shown in Fig. 4a, two exponential and a stretched exponential can fit the data well. The first exponential mode is diffusive, which is the gel mode. The second exponential term is non-diffusive, indicate the non-ergodic heterogeneity of the gel. The stretched exponential is also non-diffusive, with β=1/3. Besides, the characteristic decay time of the non-diffusive second exponential mode is 77.25ms, which is slower than the characteristic decay time of the non-diffusive stretched exponential mode, which is 3.19ms (shown in Fig. 4a)

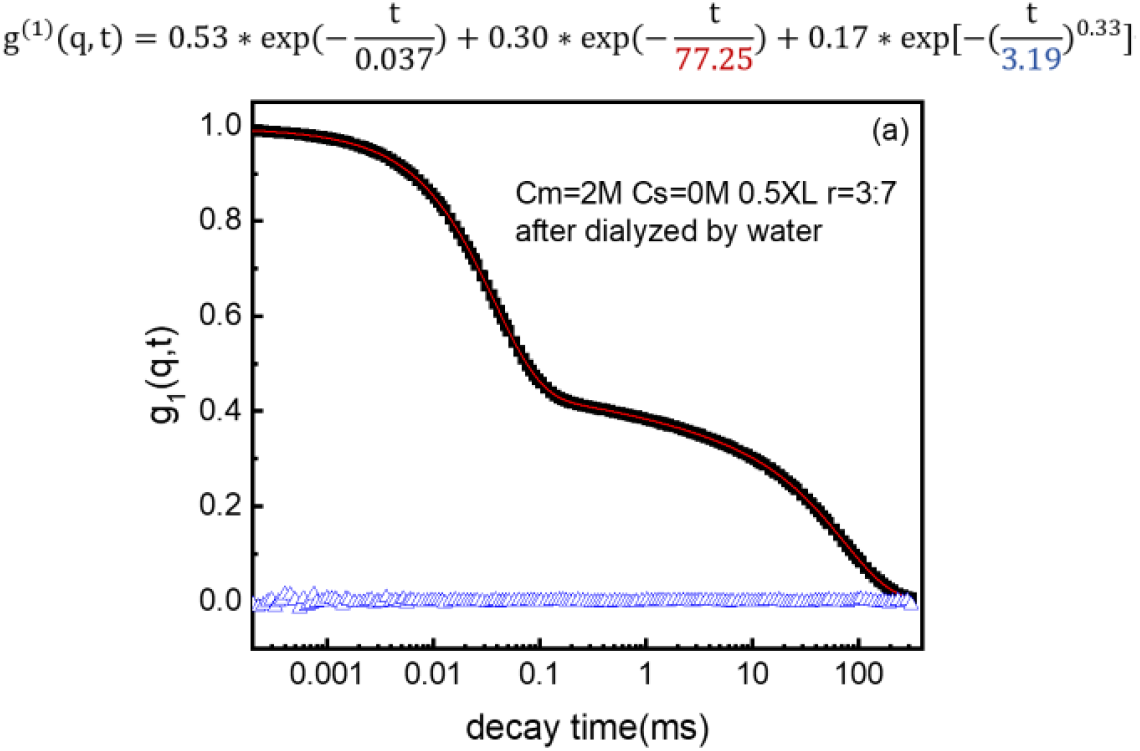

#### 4.2 Gel with molar charge ratio r of negatively charged monomers/ positively charged monomers=7:3

at fixed Cm=2M, Cs=0M (no addition salt), 0.5% crosslinking density, dialysis by water. As shown in Fig. 4b, two exponential and a stretched exponential can fit the data well. Both the first and second exponential modes are diffusive. The third stretched exponential is non-diffusive, with β=0.36. Besides, the slowest characteristic decay time occurs at the non-diffusive stretched exponential mode, which is 8.19 ms (shown in Fig. 4b). This is very different from the situation in Fig. 4a, where r =3:7

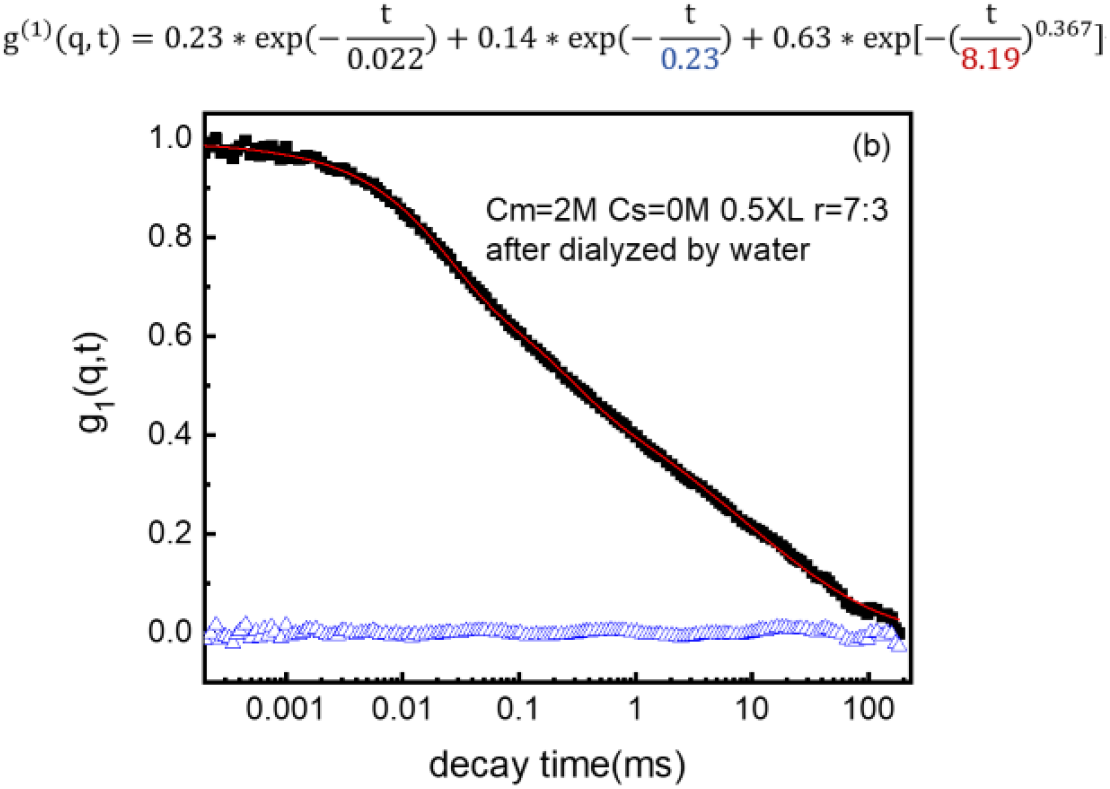

#### 4.3 gel with molar charge ratio r =1 at fixed Cm=2M, Cs=0M

(no addition salt), 0.5% crosslinking density, dialysis by water. As shown in Fig. 4c, two terms with one exponential and a stretched exponential can fit the data well. The first exponential mode is diffusive, which is the gel mode. The stretched exponential is non-diffusive, with β=0.38

### 5. 100% positively charged gel with molar charge ratio r =0 at fixed Cm=2M, Cs=0M (no addition salt), 0.5% crosslinking density of the as prepared state and dialysis by water

Surprisingly, when r=0, which means the gel is 100% positively charged gel, the stretched exponential mode disappear! For the as-prepared gel with r =0 at fixed Cm=2M, Cs=0M (no addition salt), 0.5% crosslinking density, two exponential modes can fit the data well, and both exponential modes are diffusive, with D_1_=3.19E-6(cm^2^/s), which indicates the gel elasticity, and D_2_=1.41E-7(cm^2^/s), probably arising from the coupled motion between counterions coupled with positively charged gel strands, as shown in fig. 5a-b.

**FIG. 4.**
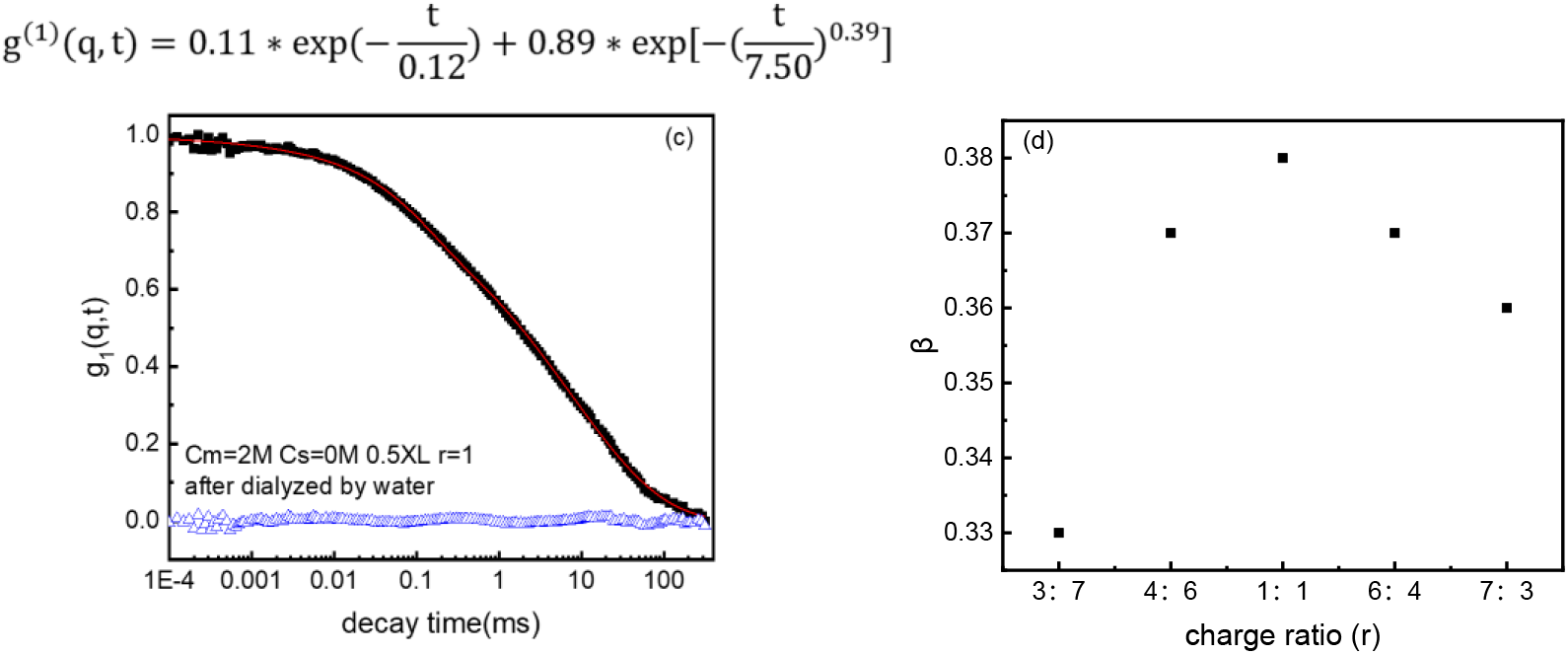
(a) Normalized field correlation function g_1_(t) at scattering angle 60° for after dialyzed gel at fixed Cm=2M, Cs=0M, 0.5XL, molar charge ratio of negatively charged monomers/ positively charged monomers r=3:7 and 0.5XL (red lines are the best fits and blue triangles are residuals). (b) Normalized field correlation function g_1_(t) at scattering angle 60° for after dialyzed gel at fixed Cm=2M, Cs=0M, molar charge ratio of negatively charged monomers/ positively charged monomers r=7:3 and 0.5XL (red lines are the best fits and blue triangles are residuals). (c) Normalized field correlation function g_1_(t) at scattering angle 60° for after dialyzed gel at fixed Cm=2M, Cs=0M, r=1 and 0.5XL (red lines are the best fits and blue triangles are residuals). (d) Stretched exponent β of after dialyzed gel at fixed Cm=2M, Cs=0M, 0.5XL with different charge ratio r.

**FIG. 5.**
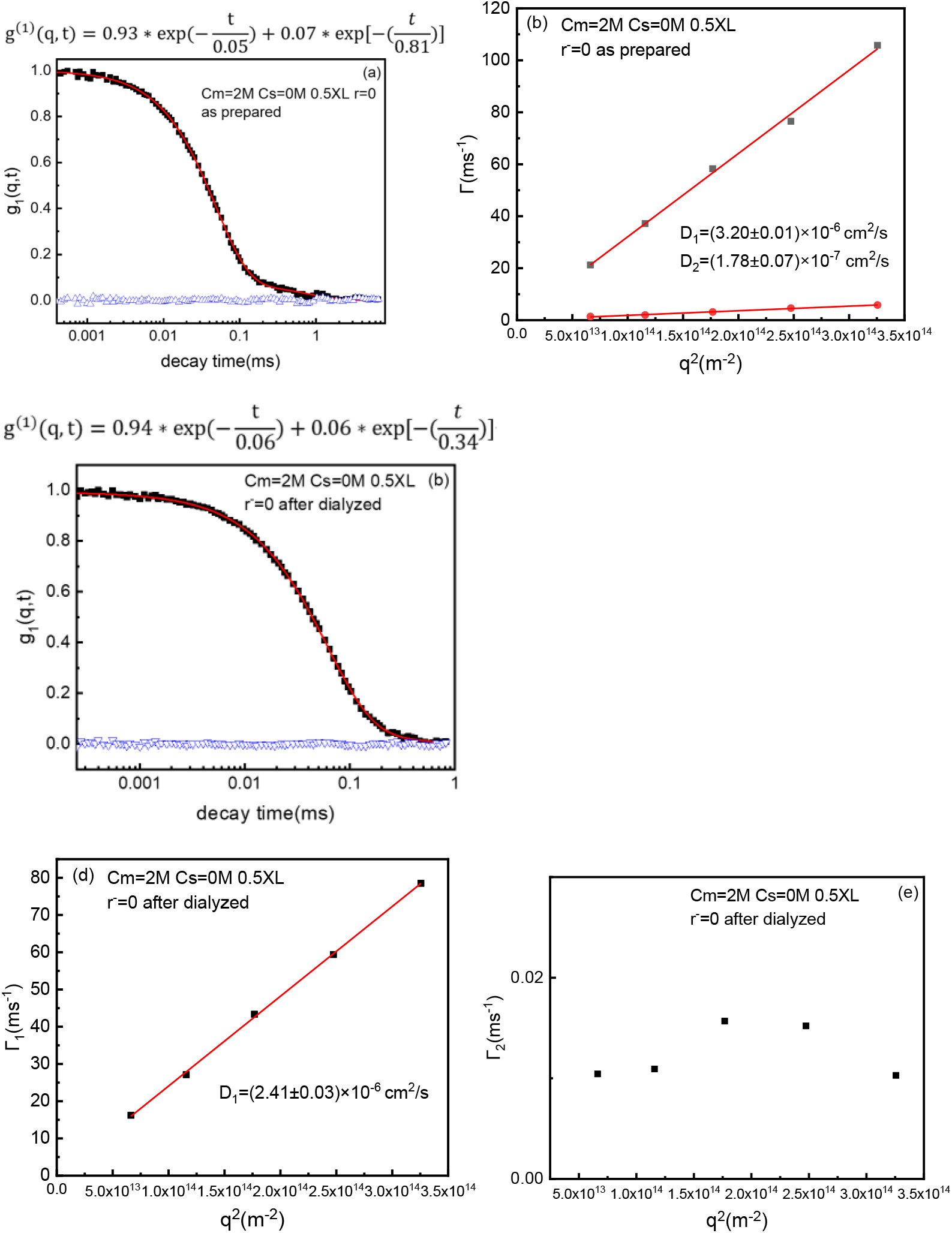
(a) Normalized field correlation function g_1_(t) at scattering angle 60° for Cm=2M, Cs=0M, 0.5XL and r=0 as prepared gel (red lines are the best fits and blue triangles are residuals). (b) Corresponding q^2^ dependence of relaxation rates Г for as prepared gel at fixed Cm=2M, Cs=0M, 0.5XL and r=0. (c) Normalized field correlation function g_1_(t) at scattering angle 60° for gel at fixed Cm=2M, Cs=0M, 0.5XL and r=0 after dialysis by water (red lines are the best fits and blue triangles are residuals). (d), (e) Corresponding q^2^ dependence of relaxation rates Г for gel at fixed Cm=2M, Cs=0M, 0.5XL and r=0 after dialysis by water.

After dialysis by water for the gel with r =0 at fixed Cm=2M, Cs=0M (no addition salt), 0.5% crosslinking density, the stretched exponential mode also disappear! And two exponential modes can also fit the data well, but only the first exponential is diffusive with D_1_=2.40E-6 (cm^2^/s). And the second exponential mode is non-diffusive, which indicates the non-ergodic heterogeneity of the gel, as shown in fig. 5c-e. This case goes back to the normal DLS results of the regular charged gels, such as Poly(acrylate-co-amide) gels. Normally, the charged gels have some heterogeneity, so that they will exhibit a non-diffusive exponential decay as a second mode.

## ACKNOWLEDGMENTS

This work was supported by the National Key R&D Program of China (Grant No. 2023YFE0124500), the National Natural Science Foundation of China (Grant No. 22273114), the National Key R&D Program of China (Grant No. 2023YFC2411203), and International Partnership Program of the Chinese Academy of Sciences (Grant No. 027GJHZ2022061FN).

## Notes

### Competing Interest Statement

The authors have declared no competing interest.

