## Supplemental Figure 1 for "Interplay of hierarchical dynamics and their microscopic structures of polyampholyte gels and proteins"

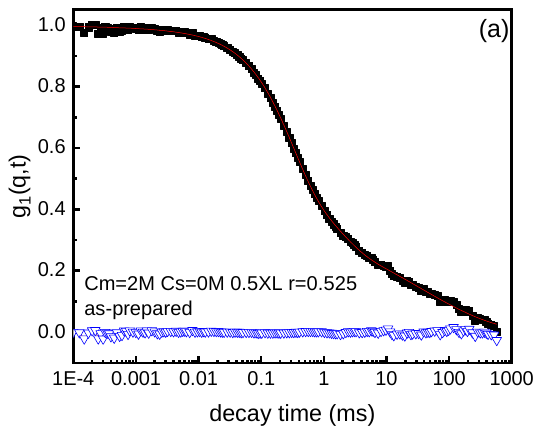

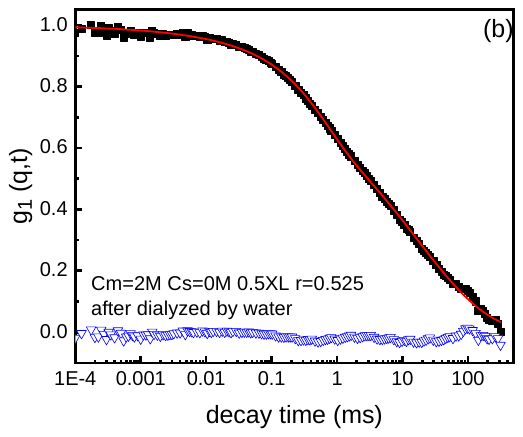

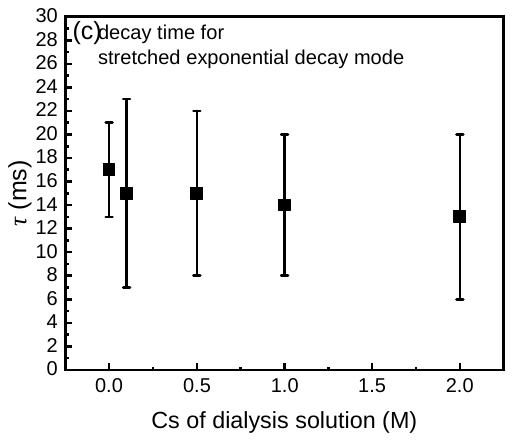

**Fig. 1 a,** Normalized field correlation function g_1_(t) at scattering angle 30° for as-prepared gels at fixed Cm=2M, Cs=0M, r=0.525 and 0.5XL (red lines are the best fits and blue triangles are residuals). **b,** Normalized field correlation function g_1_(t) at scattering angle 30° for gels after dialyzed by water at fixed Cm=2M, Cs=0M, r=0.525 and 0.5XL (red lines are the best fits and blue triangles are residuals). **c,** Decay time for stretched exponential decay of Cm=2M, Cs=0M, r=0.525, and 0.5XL gels at different Cs of dialysis solution at scattering angle 60°.
